# Development of an adenosquamous carcinoma histopathology-selective lung cancer graft model

**DOI:** 10.1101/2022.08.23.504928

**Authors:** I.A.K. Lähdeniemi, J.R. Devlin, A.S. Nagaraj, S.S. Talwelkar, J. Bao, N. Linnavirta, C. Şeref Vujaklija, E.A. Kiss, A. Hemmes, E.W. Verschuren

## Abstract

Preclinical tumor models with native tissue microenvironments provide essential tools to understand how heterogeneous tumor phenotypes relate to drug response. Here, we present syngeneic graft models of aggressive, metastasis-prone histopathology-specific NSCLC tumor types driven by *KRAS* mutation and loss of *LKB1* (KL): adenosquamous carcinoma (ASC) and adenocarcinoma (AC). We show that subcutaneous injection of primary KL-ASC cells results in squamous cell carcinoma (SCC) tumors with high levels of stromal infiltrates, lacking the source heterogeneous histotype. Despite forming subcutaneous tumors, intravenously injected KL-AC cells were unable to form lung tumors. In contrast, intravenous injection of KL-ASC cells leads to their lung re-colonization and lesions recapitulating the mixed AC and SCC histopathology, tumor immune suppressive microenvironment and oncogenic signaling profile of source tumors, demonstrating histopathology-selective phenotypic dominance over genetic drivers. Pan-ERBB inhibition increased survival, while selective ERBB1/EGFR inhibition did not, suggesting a role of ERBB network crosstalk in resistance to ERBB1/EGFR. This immunocompetent NSCLC lung colonization model hence phenocopies key properties of the metastasis-prone ASC histopathology, and serves as a preclinical model to dissect therapy responses and metastasis-associated processes.

## Introduction

Non-small cell lung cancer (NSCLC) is the most common form of lung cancer (85%) and one of the leading cancer-related causes of deaths in the world. NSCLC is a heterogeneous disease that is often discovered at late stages resulting in poor prognoses and clinical outcomes (Bray et al., 2018). Novel tyrosine kinase inhibitors (TKI) and immunotherapy agents have provided significant clinical promise for NSCLC treatment in the past decade with approved TKIs such as erlotinib and afatinib demonstrating increased progression-free survival compared to conventional chemotherapy (Liu et al., 2021). Immunotherapy with programmed cell death 1 (PD-L1) checkpoint inhibitors is approved for metastatic NSCLC treatment and significantly increases patient survival rates (Brahmer et al., 2015). However, heterogeneity of immune checkpoint protein expression is a feature of NSCLC that limits the broad effectiveness of these therapies (Gettinger et al., 2015), with the absence of PD-L1 expression identified in both human and murine tumors with oncogenic *KRAS* and loss-of-function *LKB1* mutations (Calles et al., 2015, Koyama et al., 2016, Skoulidis et al., 2015). The absence of checkpoint marker expression in pre-clinical models of dual *KRAS; LKB1* mutated NSCLC corresponded with poor to no-responsiveness to checkpoint immunotherapy and subsequently effective treatments to durably target tumors that are non-responsive to immunotherapy or TKIs remain lacking (Skoulidis et al., 2018, Li et al., 2022).

Genetically engineered mouse models (GEMMs) constitute excellent tools to model the diversity of NSCLC biology, allowing assessment of therapeutic responses within the context of native immunocompetent microenvironments. Our previous findings highlight the importance of the cell-of-origin in histotype-selective growth of tumors driven by oncogenic *Kras*^G12D^ mutation and loss of *Lkb1* (KL). Tumors initiated in club cell antigen 10 (CC10) progenitors led to predominant formation of mixed histopathology adenosquamous carcinoma (ASC) tumors and a wider range of histotypes compared to KL tumors originating from surfactant protein C (SPC) progenitors that predominantly exhibited adenocarcinoma (AC) histopathology (Nagaraj et al., 2017). Interestingly, KL;ASC tumors exhibited a histotype-selective immune suppressive microenvironment with decreased expression of MHC genes and decreased T-cell infiltration, as well as increased recruitment of CD11b^+^ GR1^+^tumor-associated-neutrophils (TANs) (Nagaraj et al., 2017, Schabath et al., 2016).

GEMMs have served as a great tool to study cancer biology including localized progression and distal metastasis, which are both influenced by multiple factors including the immune system, TME and driver genes. GEMM studies have contributed to development of novel promising therapeutics which have been shown to be effective for specific subsets of NSCLC such as TKI, as well as other targeted therapies and immunotherapies for other solid cancer types (Yuan et al., 2019). Despite their advantages to dissect disease pathology, the extended time frames for tumor development as well as intra- and inter-cohort heterogeneity limits the utility of GEMMs for accelerated evaluation of anti-cancer drug efficacy. While advances have been made for the establishment of faithful *in vitro* and *ex vivo* approaches to model both patient- and GEMM-derived tumors, these systems exhibit limitations in their ability to fully recapitulate the complexities of *in vivo* tumor biology and consequently to optimally reflect therapeutic responses (Chaffer and Weinberg, 2011, Quinn et al., 2021).

To address methodological limitations in evaluation of cancer treatment approaches, we and others have developed and validated methods for NSCLC functional diagnostics, including conditionally reprogrammed cells (CRCs) (Liu et al., 2012), long-term 3D organoid culture (Kim et al., 2019, Sachs et al., 2019) and analysis of functional uncultured tumor cells (FUTCs) (Talwelkar et al., 2021). These *ex vivo* models can complement GEMMs in preclinical cancer research. For example, culture and analysis of KL GEMM-derived primary cells specifically uncovered NSCLC histotype-selective drug sensitivity and bypass resistance mechanism. MEK inhibitor (trametinib) treatment was shown to be selective for AC tumors, while both ASC and AC histotypes showed response to the pan-ERBB inhibitor afatinib. *Ex vivo* data was validated *in vivo*, showing afatinib response in both KL;ASC and KL;AC tumors (Talwelkar et al., 2019).

GEMMs exhibit limitations as preclinical models due to relatively long tumor development times as well as inter-tumor histotype diversity between mice within treatment cohorts. We here set out to develop and compare subcutaneous (s.c.) and intravenous (i.v.) transplantation models to study KL-ASC and -AC histotype tumors in both immune-competent and -compromised hosts. We observed differential metastatic capacity of histotype-selective cells, unexpectedly showing that only ASC cells were capable of re-colonizing the lung after i.v. transplantation. Importantly, these lung colonized tumors retained the key mixed AC and SCC histotype features as well as immunosuppressive tissue microenvironment (TME) of source ASC tumors, which was not seen with the s.c. model. We analyzed responses to inhibitors of ERBB family receptor tyrosine kinases (RTKs,) that have been previously shown to drive tumor progression in the KL GEMM, and found that only pan-ERBB inhibition increased the survival of ASC^i.v.^ mice compared to control or single ERBB1/EGFR inhibition. Taken together, this syngeneic ASC^i.v.^ transplantation model will be useful to evaluate therapy responses in microenvironments that accurately recapitulate GEMM biology, and will help to analyze the mechanisms behind the differential metastatic capacity of ASC tumor cells.

## Results

### Development of a NSCLC metastatic lung cancer model

Our previous studies in mice have revealed that selective expression of the oncogenic *Kras*^G12D^ allele in combination with *Lkb1* deletion (KL) in lung CC10 cells leads to formation of aggressive ASC lesions, in addition to other tumor histotypes, while ASC histopathology was not observed in tumors arising from SPC cells harboring the same genetic mutations (Nagaraj et al., 2017). To investigate the histopathology-selective phenotypes of NSCLC cancer, including potential differences in their transplantation capacity *in vivo*, we first assessed the tumor-forming capacity of bulk cell populations isolated from GEMM-derived KL;ASC and KL;AC cells outside of the lung. AC^s.c^ and ASC^s.c.^ were generated by subcutaneously injecting KL;ASC and KL;AC cells into the flanks of immune compromised athymic nude mice (Figure 1A). Tumors derived from KL;ASC transplanted cells (ASC^s.c^) exhibited accelerated growth compared to those derived from KL;AC cells (AC^s.c^) (Figure 1B), consistent with the greater proliferative capacity of GEMM-derived KL;ASC cells that has been previously reported *ex vivo* (Talwelkar et al., 2019). This could also be seen *in vivo* where ASC^s.c.^tumors displayed high Ki-67 positivity compared to AC^s.c.^ tumors (Figure 1C and S1B). This phenotype was recapitulated using KL;ASC and KL;AC cells which had been established as *ex vivo* cultures prior to s.c. transplantation (Figure S1A). Interestingly, immunohistochemical (IHC) end-point analyses revealed that independent of the tumor source (KL;ASC vs KL;AC), resulting s.c. tumors did not exhibit staining for the AC transcription factor 1 (nkx2-1: Figure 1C). ASC^s.c.^ lesions exhibited strong staining for the squamous cell (SCC) marker p63, which was maintained from the source KL;ASC tumor (Figure 1C) while positive p63 staining in AC^s.c.^ tumors appeared to be associated with stromal cell infiltration rather than a transition to squamous cell histopathology. Stromal cell infiltration was confirmed with vimentin staining which indeed showed high positivity in both s.c. tumor types (Figure 1C). Loss of nkx2-1 staining was also observed in ASC^s.c.^ and AC^s.c.^tumors grown in immune-competent C57BL/6 mice following transplantation of background-matched KL;ASC and KL;AC bulk cell populations (Figure 1D). Consistent with the immune-compromised mice, these tumors exhibited strong positivity for the mesenchymal marker vimentin, further suggesting stromal cell infiltration. Ki-67 stained positive for a subset of cells suggesting maintenance of proliferative capacity and no differences could be observed between the lesions established in both host strains (Figure 1D, S1B). The loss of key histopathological markers and stromal cell infiltration suggests that subcutaneous transplantation approach will not be the best system to model the complexity of NSCLC subtypes for the evaluation of anti-cancer therapy responses.

**Figure 1:**
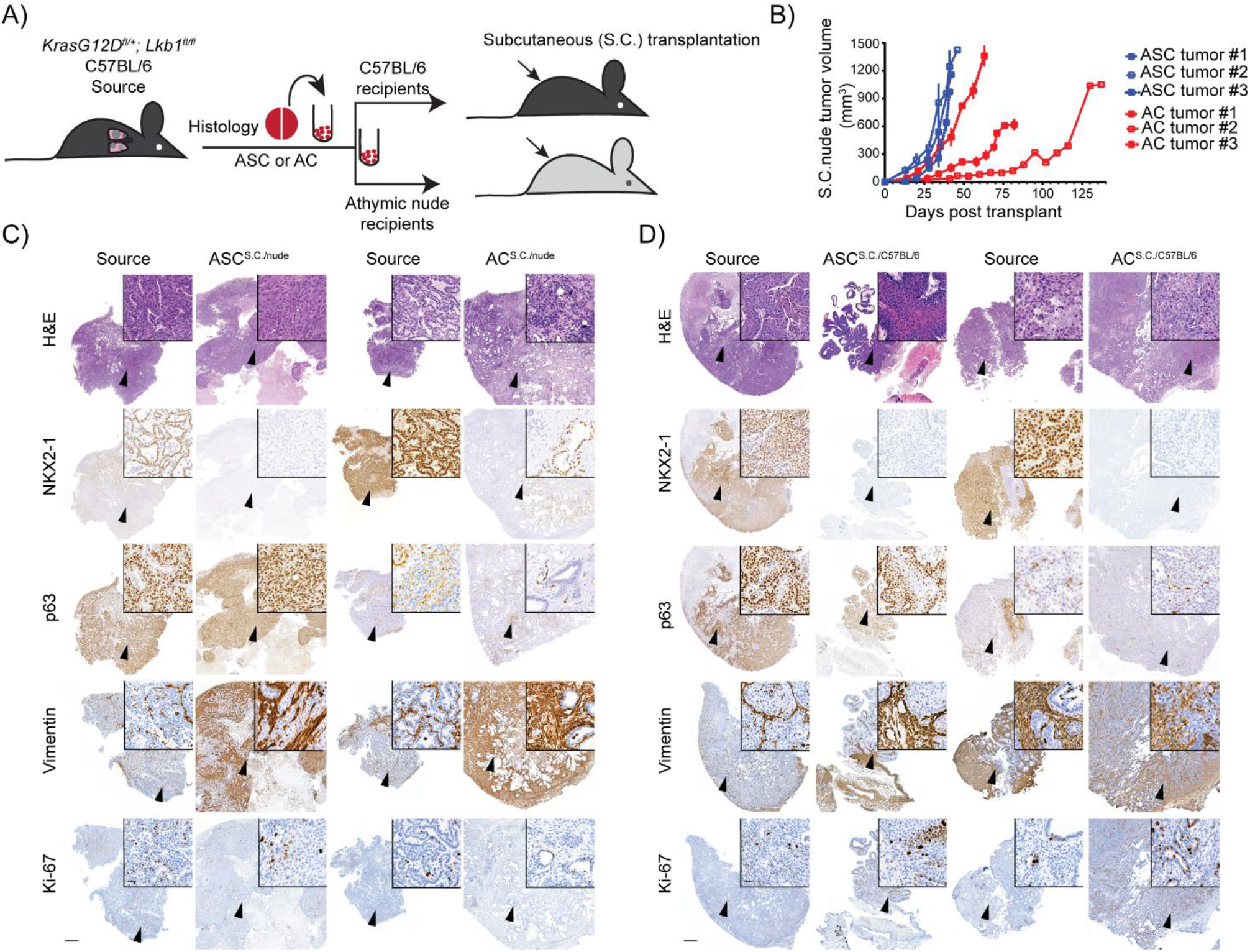
Host microenvironment-dependent and histopathology-selective tumor establishment subcutaneously. **(A)** Schematic overview of generation of syngeneic or immune compromised graft models with subcutaneous (s.c.) transplantation of ASC or AC tumor-derived cells from KL nude or C57BL/6 mice. Single cell suspensions derived from dissociated KL C57BL/6 source tumors were subjected to dead cell removal prior to transplantation into syngeneic or immune-compromised recipients. A reference tissue piece was fixed from tumors for H&E and IHC staining and confirmation of tumor histopathology. (**B**) Subcutaneous ASC or AC tumor growth curves in athymic nude recipient mice. Each line represents one mouse and the tumors were measured weekly by using a caliper. n=3. (**C)** Representative H&E, nkx2-1/ttf1 and p63 IHC images of source and s.c. tumors from athymic nude recipient mice upon transplantation with ASC or AC tumor-derived cells. Scale bar 500 μm (2x) or 20 μm (40x) n=3. (**D)** Representative H&E, nkx2-1 and p63 IHC images of source and s.c. tumors from C57BL/6 recipient mice upon transplantation with ASC or AC tumor-derived cells. Scale bar 500 μm (2x) or 20 μm (40x) n=6.

Metastatic models where breast cancer cells have been intravenously (i.v.) transplanted via mouse tail vein injection have been used to study the role of immune cells in modulating the metastatic processes (Li et al., 2020, Tyagi et al., 2021). To study the metastatic capability of histopathology-selective NSCLC cells, bulk cell populations from KL;ASC and KL;AC GEMM tumors were i.v. transplanted into athymic nude and C57BL/6 mice (Figure 2A). In contrast to our observations following s.c. transplantation, only cells from KL;ASC. tumors exhibited the capacity to recolonize the lung, while no tumor formation was observed following i.v. transplantation of KL;AC cells, independent of host immune-competence (Figure 2B-F). ASC^i.v.^ tumors were variable in number and size, with a median of 9 tumors per lung counted in nude hosts and 19.9 tumors per lung measured in C57BL/6 hosts (Figure 2B-D, S2A and B). Post-mortal IHC analysis revealed that, again in contrast to results observed in ASC^s.c.^ (Figure 1C), both AC (nkx2-1) and SCC (p63) characteristics were maintained in ASC^i.v.^ tumors, with the transplanted lesions exhibiting comparable histopathology to the source tumors independent of the host (Figure 1D and E, S2A and B). KL tumors have an adaptive activation of receptor tyrosine kinase (RTK) signaling pathways such as the ERBB, MET and MAPK signaling cascades (Manchado et al., 2016). ASC^i.v.^ tumors exhibited positive staining for phosphorylated AKT (pAKT) and ERK1/2 (pERK) that was comparable to the ASC source tumors (Figure 2E and F; Figure S3A and B). Consistent with our previous report (Narhi et al., 2018), AC source tumors displayed significantly decreased pAKT levels compared to both ASC^i.v.^ and ASC source tumors, and instead exhibited stronger pERK staining (Figure S3A and B). While no differences in pEGFR levels could be observed between different mice, for ASC comparable levels of phosphorylated ERBB2 and ERBB3 were observed between source and the transplant-derived tumors, with significantly lower pERBB2 and pERBB3 levels detected in AC source tumors (Figure S3C-F) consistent with our previous observations (Talwelkar et al., 2019). Consequently, in addition to the maintenance of histopathological features, oncogenic signaling pathway activity is maintained in intravenously transplanted ASC lesions compared with source GEMM tumors (Figure S3C-F).

**Figure 2:**
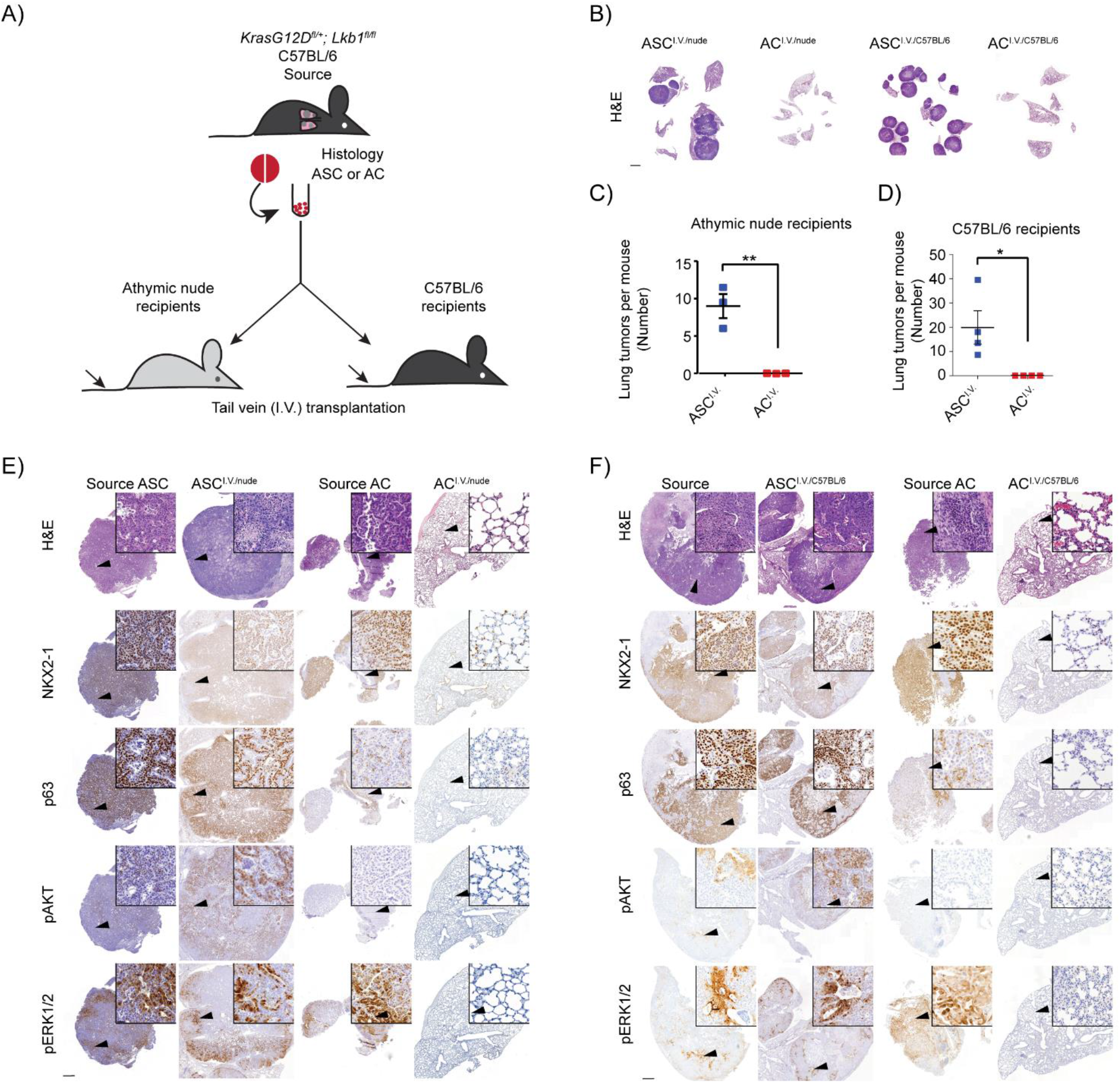
KL;ASC cells re-colonize to the lung and form tumors after intravenous injection while AC KL deficient cells form no tumors. **(A)** Schematic overview of generation of syngeneic or immune-compromised graft models with intravenous (i.v.) transplantation of ASC or AC tumor-derived cells from KL athymic nude or C57BL/6 mice. Single cell suspensions derived from dissociated KL C57BL/6 source tumors were subjected to dead cell removal prior to transplantation into syngeneic or immune-compromised recipients. A reference tissue piece was fixed from tumors for H&E and IHC staining and confirmation of tumor histopathology. **(B)** Representative images of nude and C57BL/6 ASC^i.v.^ and AC^i.v.^ mice whole lungs stained for H&E. Scale bar 2000 μm (0.4x). n=6. **(C-D)** The number of tumors / mouse was counted based on the H&E stainings done on tumor tissues. The graph represents mean +/− SD for each group, and each point represents an individual mouse. Two-tailed unpaired Student’s t-test values are *p<0.05, **p<0.01. n=3 (nude); 4 (C57BL/6). **(E)** Representative H&E, ttf1, p63, pAKT and pERK1/2 IHC images of source and i.v. tumors from athymic nude recipient mice upon transplantation with ASC or AC tumor-derived cells. Scale bar 500 μm (2x) or 20 μm (40x) (n=3) (**F)** Representative H&E, Nkx2-1, p63, pAKT and pERK1/2 IHC images of source and ASC^i.v.^ tumors from C57BL/6 recipient mice upon transplantation with ASC or AC tumor-derived cells. Scale bar 500 μm (2x) or 20 μm (40x). n=4.

### Maintenance of source tumor-intrinsic and host-systemic immunophenotype of i.v. transplanted ASC lesions

Our previous data indicated a neutrophil-dominating, low T-cell immune phenotype in KL;ASC GEMM tumors (Nagaraj et al., 2017). We therefore assessed whether the immune phenotype of our ASC^i.v.^ tumors recapitulated that of the source tumor using IHC analysis of the CD11b neutrophil marker and CD3 T-cell marker. CD3 positivity remained mainly negative throughout ASC^i.v^ tumors in all mice while CD11b staining indicative of neutrophil infiltration was more prominent and consistent between ASC^i.v^ lesions and GEMM source tumors (Figure 3A-B). There were no differences in the percentage of CD11b positive cells between source KL;ASC and ASC^i.v.^ tumors (Figure S4).

**Figure 3.**
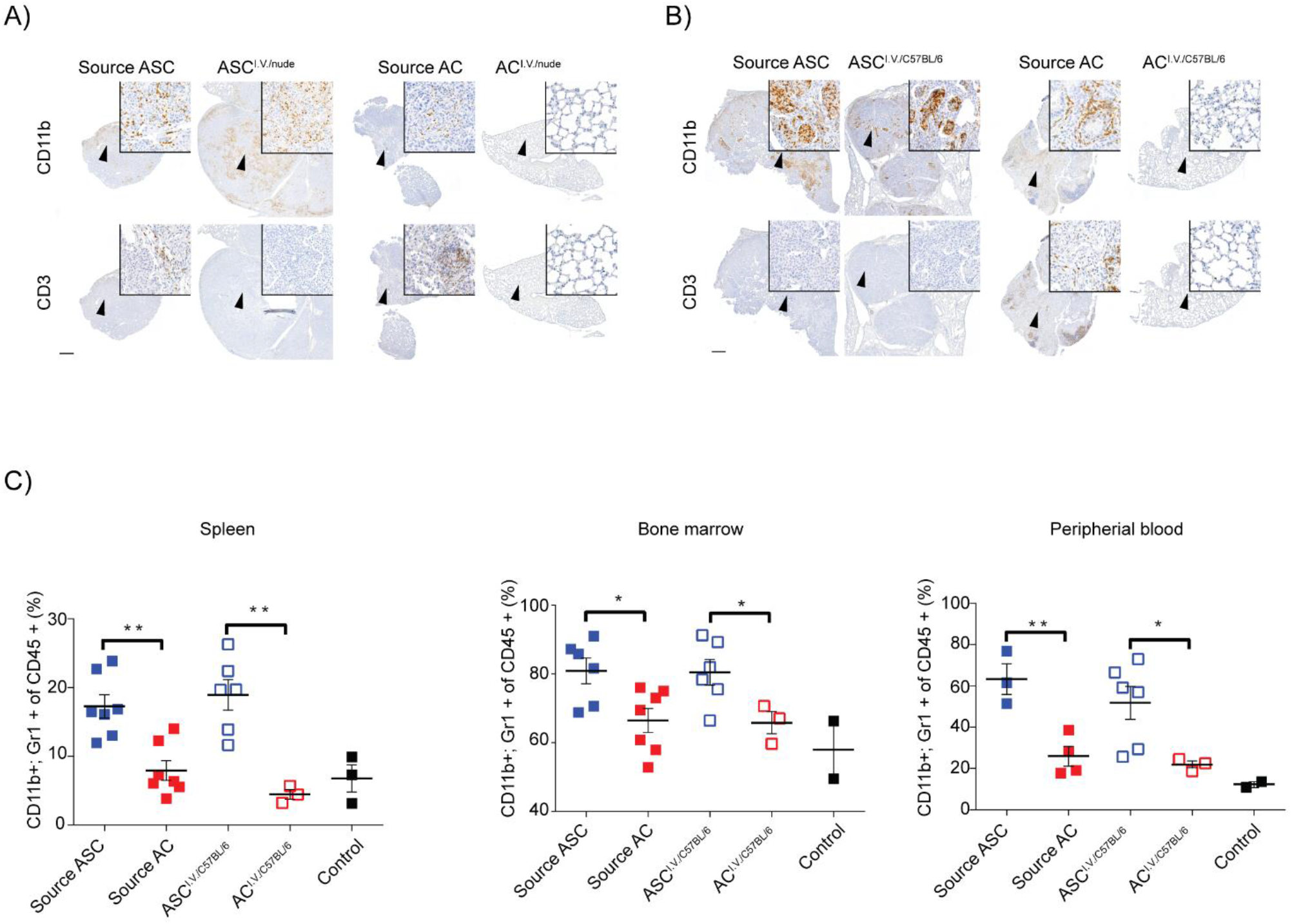
The immune phenotype in the ASC^i.v.^ model resembles the source ASC immune phenotype. **(A)** Representative CD11b and CD3 IHC images of source and i.v. tumors from athymic nude recipient mice upon transplantation with ASC or AC tumor-derived cells. Scale bar 500 μm (2x) or 20 μm (40x). n=3. **(B)** Representative CD11b and CD3 IHC images of source and i.v. tumors from C57BL/6 recipient mice upon transplantation with ASC or AC tumor-derived cells. Scale bar 500 μm (2x) or 20 μm (40x). n=3. **(C)** Flow cytometric analysis of CD11b+;Gr1+ cells (as a percentage of total CD45+ leukocytes) in bone marrow, peripheral blood and dissociated spleens isolated from ASC-source (Ad5-CC10-Cre infected KL mice), or C57BL/6 recipient mice i.v. transplanted with ASC tumor-derived cells. Graph represents the mean +/− SEM of each group and each point represents an individual mouse. Two-tailed unpaired Student’s t-test values are *p<0.05, **p<0.01. n=7.

In addition to assessment of tumor-intrinsic immune phenotypes, the relationship between tumor histopathology and the host’s systemic immune profile was assessed in the C57BL/6 background. Using flow cytometry, CD11b, GR1 positive cells were analyzed in bone marrow, spleen and peripheral blood in both the GEMMs from which source tumors were originally isolated and the host mice which were subsequently transplanted. GEMMs with KL;ASC tumors had significantly higher levels of CD11b positive cells in all three compartments compared to GEMMs with KL;AC tumors which had similar systemic neutrophil populations to non-tumor bearing control mice (Figure 3C). These results complement previous findings where KL;ASC tumors were reported to have an increased TAN recruitment compared to KL;AC tumors (Nagaraj et al., 2017). Importantly, host mice with ASC tumors also exhibited significantly elevated CD11b, GR1 positive cell populations in the spleen, bone marrow and peripheral blood compared to mice transplanted with KL;AC cells (Figure 3C). Taken together, the histopathological and immune profiling studies indicate that i.v. transplantation of KL;ASC cells into C57BL/6 hosts (ASC^i.v^.^/C57BL/6^) reliably recapitulates the source GEMM tumor phenotype, rendering it suitable to evaluate anti-cancer treatments for this aggressive subtype of NSCLC.

### Therapeutic responses in the ASC^i.v.^ lung transplantation model recapitulate KL;ASC tumors

To investigate how well the i.v. transplantation model mimics NSCLC responses to previously established therapies, we treated ASC^i.v./C57BL/6^ mice with the pan-ERBB inhibitor afatinib (AF). Drug treatments commenced 10 days post-transplantation and continued for four weeks, at which point lung tumor burden was analyzed. Mice in the vehicle control group developed large ASC tumors with 75.5% lung area defined as tumor, while AF-treated mice exhibited significantly lower tumor burden (15.7%; Figure 4A and B), consistent with previously observed therapeutic response in KL-ASC GEMMs (Talwelkar et al., 2019).

**Figure 4:**
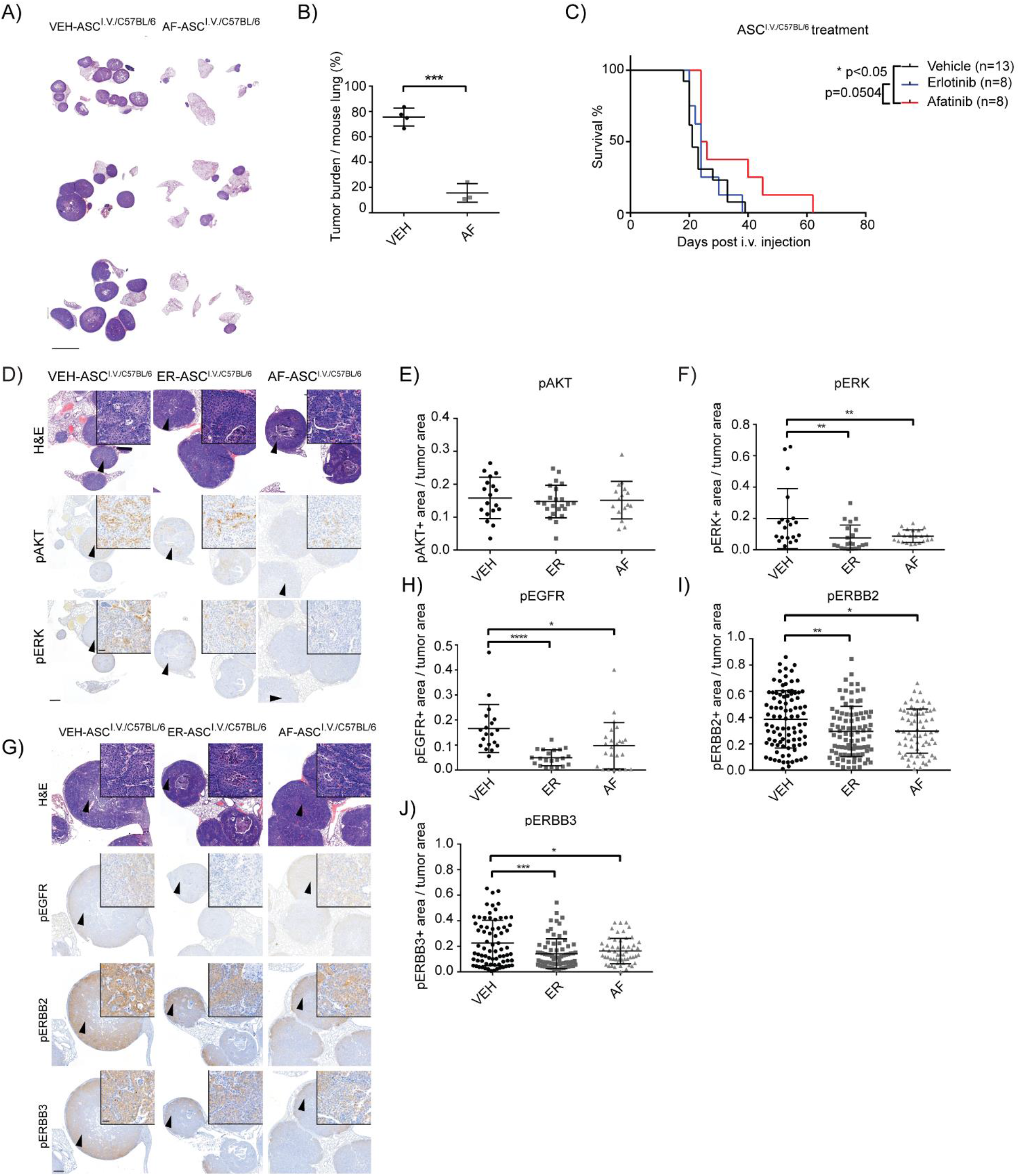
ERBB inhibition with afatinib and erlotinib decreases the number of ERBB and pERK positive cells and pan-ERBB inhibition increases the survival of mice. **(A)** Three representative H&E stainings of mice treated for four weeks 10 days after ASC tumor cell i.v. transplantation with either vehicle control or afatinib (AF). Scale bar 5000 μm. n=4 (VEH); 3 (AF). **(B)** Quantification of the tumor burden after treatments presented in (A). The graph represents mean+/− SD for each group, and each point represents an individual mouse. Two-tailed unpaired Student’s t-test value is ***p<0.001. n=4 (VEH); 3 (AF). **(C)** The survival percentage of mice treated with vehicle, ER and AF 10 days after ASC tumor cell i.v. transplantation. Graph represents the mean +/− SD for each group, and each point represents an individual mouse. Gehan-Breslow-Wilcoxon test values are *p<0.05. n=11 (VEH); 8 (ER/AF). **(D)** Representative H&E, pAKT and pERK1/2 IHC images of source and i.v. tumors from C57BL/6 recipients upon transplantation with ASC tumor derived cells and treated with vehicle, EGFR inhibitor erlotinib (ER) or pan-ERBB inhibitor afatinib (AF). Scale bar 500 μm (2x) or 20 μm (40x). n=5. **(E-F)** Quantification of pAKT and pERK1/2 positive area by dividing the positive tumor area with the total tumor area in each tumor separately by using FIJI ImageJ. The graph represents mean+/− SD for each group, and each point represents an individual tumor. Two-tailed unpaired Student’s t-test values are **p<0.01. n=5. **(G)** Representative H&E, pEGFR, pERBB2 and pERBB3 IHC images of source and i.v. tumors from C57BL6 recipients upon transplantation with ASC tumor derived cells and treated with vehicle, EGFR inhibitor erlotinib (ER) or pan-ERBB inhibitor afatinib (AF) and sacrificed when mice developed health problems. Scale bar 500 μm (2x) or 20 μm (40x). n=5. **(H-J)** Quantification of pEGFR, pERBB2 and pERBB3 positive area by dividing the positive tumor area with the total tumor area in each tumor separately by using FIJI ImageJ. The graph represents mean+/− SD for each group, and each point represents an individual tumor. Two-tailed unpaired Student’s t-test values are *p<0.05, **p<0.01, ***p<0.001. n=5.

To further explore both the therapeutic response of transplanted tumor cells following lung colonization and the corresponding impact on host survival, ASC^i.v^.^/C57BL/6^, mice were treated with vehicle, AF or the EGFR inhibitor erlotinib (ER) from 10 days post transplantation and sacrificed when they presented with physical signs of disease burden or exhibited 20 percent body weight loss. All mice, independent of treatment group, developed ASC histotype tumors as indicated by positive nkx2-1 and p63 staining (Figure S5A). Afatinib, but not ER, treatment provided a significant survival advantage compared to vehicle control (Figure 4C), suggesting that a selective EGFR inhibitor is insufficient to provide a survival extension in this model. At end-point there was no difference between treatment groups with respect to the proportion of CD11b positive infiltrating neutrophils (Figure S5B and C), the tumor number per mouse (Figure S5D) or tumor area in the lungs (Figure S5E). However, the treatment modulated the KRAS downstream signaling pathway, with significantly decreased staining for phosphorylated ERK1/2, but not phosphorylated AKT, detected for both AF and ER treatments (Figure 4D-F). AF and ER treatment also modified the phosphorylation of key ERBBs including EGFR, ERBB2 and ERBB3, which all exhibited significantly decreased phosphorylation in both AF- and ER-treated mice (Figure 4G-J). However, for both AF- and ER-treated mice end-point tumors had populations of cells that maintained ERBB phosphorylation, with higher levels maintained in ER-treated mice. This is in line with already published data on KL mice phenotype indicating an improved therapeutic response with pan-ERBB inhibitors compared to EGFR inhibitors (Kruspig et al., 2018, Moll et al., 2018, Talwelkar et al., 2019) further suggesting that our i.v. transplantation model is a reliable preclinical tool for NSCLC therapy and metastasis analysis.

## Discussion

There is a need to develop improved, tractable *in vivo* models of NSCLC that accurately reflect the original tumor histopathology, oncogenic signaling profile and immunogenic landscape that can be used to assess therapeutic treatments and potentially provide insights about NSCLC metastatic processes. NSCLC is a problematic disease to diagnose and is often diagnosed at a late stage when the disease has already developed metastatic expansions. Even though the therapeutics of NSCLC has developed enormously during recent years, a deeper understanding about NSCLC signaling profiles and metastasis processes is still necessary (Kim et al., 2020). We have here created a reproducible and rapid tumor-lung colonization model to study an aggressive subset of NSCLC via i.v. transplantation of GEMM-derived tumor cells into healthy recipient mice. Lung-colonizing ASC tumors maintained a previously characterized histopathology-associated oncogenic signaling phenotype displayed in the source GEMM, and in immune-competent C57BL/6 hosts exhibited a previously identified immune-suppressive phenotype characterized by TAN infiltration. Moreover, an unappreciated systemic immune landscape associated with ASC but not AC NSCLC histopathology, and characterized by the expansion of CD11b; Gr1 positive neutrophils in the spleen, bone marrow and bloodstream, was maintained between GEMM and i.v. transplanted mice. The i.v. transplantation model accurately recapitulated ASC sensitivity to the clinically relevant pan-ERBB inhibitor afatinib, through both short-term ablation of localized tumor progression and extended survival upon longer treatment (Moll et al., 2018, Talwelkar et al., 2019).

Despite the equivalent ability of cells derived from GEMM ASC- and AC-lesions to form tumors upon subcutaneous (s.c.) transplantation of both immune-compromised and -competent hosts, lung-colonization upon i.v. transplantation was limited to ASC cells with AC-injected mice exhibiting no lung tumors or any other indication of disease burden. ASC lesions have been reported to be more aggressive and have a poor prognosis when compared to AC tumors (Nagaraj et al., 2017, Pan et al., 2018) and potential molecular mechanisms underpinning differential colonization / metastatic capacity of tumor cells harboring equivalent genetic driver mutations (*Kras*^*G12D*/+^; *Lkb1*^-/-^) but diverse histopathology are intriguing. One of the threshold elements in the metastatic process is the breakage of the extracellular matrix (ECM) and initiation of EMT, which associated with drug resistance and is typified by the loss of e-cadherin and cell-to-cell connections in favor of increased expression of mesenchymal markers such as expression of vimentin (Peinado et al., 2007, Singh and Settleman, 2010). ASC tumors have been reported to have high levels of chemokine receptors that are associated with increased invasiveness and chemoresistance through insensitivity to apoptotic induction (Zhu et al., 2020). This has been proposed to be dependent upon altered oxidative signaling; oxidative stress has been reported to inhibit distant metastasis of human melanoma cells in mice (Piskounova et al., 2015). Further studies will be required to determine whether this is a potential route through which ASC cells exhibit superior lung colonization capacity compared to ACs.

In addition to tumor cell phenotypes and genetic drivers, the microenvironment of the tumor plays an important role for both local tumor development as well as metastasis. Interestingly, the maintenance of tumor histopathology was compromised by localized s.c. injection of the ASC and AC cells, with transplant-derived lesions exhibiting stromal infiltration and loss of the AC marker nkx2-1. In contrast both AC and SCC features were maintained in lung-colonizing tumors formed following ASC i.v. transplantation pointing to the potential importance of the lung microenvironment for development of key histopathological features that have previously predicted response to therapy. Thus this i.v. lung colonization model serves as a promising tool to investigate the genetic and potentially non-genetic drivers of NSCLC metastases in a histopathology-selective way, and to assess their response to different treatment options.

Adaptive responses driving resistance to targeted drug therapies in tumors is a severe problem and has in the NSCLC field promoted the preclinical exploration of novel treatment options such as combinatorial treatments with MEKi and EGFRi (Kitai et al., 2016). Several reports promote the rationale for combination treatment approaches (Kruspig et al., 2018, Manchado et al., 2016), however we have previously investigated the responses of KL GEMMs to afatinib and trametinib treatment and found that combinatorial treatment does not significantly extend survival beyond single-agent treatments (Talwelkar et al., 2019). Whether this conflicting observation is a consequence of intra- and inter-mouse tumor heterogeneity in the KL GEMMs that is absent in cell-line based xenograft therapy studies is an open question. Thus, we sought to determine if our GEMM-derived lung colonization model could serve as an alternative tool to model *KARS*-mutant NSCLC responses to novel therapeutic strategies. The i.v. ASC transplantation model robustly recapitulated our previously reported results using KL GEMMs and *ex vivo* culture systems with AF single treatment dramatically ablating tumor growth in the lungs upon short-term treatment and significantly prolonging survival following extended therapy, a benefit not provided by single-agent ER treatment which is consistent with earlier studies (Kruspig et al., 2018, Moll et al., 2018). Profiling of phosphorylated ERBB2 and ERBB3 in end-stage tumors indicated that on-target activity of AF was sustained despite tumor progression, pointing to the emergence of alternative resistance mechanisms which could be explored further using this transplantable model system. Investigating the i.v. transplanted ASC tumor cells with lineage tracing could provide answers on what adaptive signaling pathways are activated upon treatments and if combinatorial therapy could provide an increased efficacy.

In addition to agents targeting key NSCLC oncogenic pathways, our lung colonization model also provides opportunities to assess other therapeutic strategies including immune-focused approaches. Neutrophils often predominate the immune landscape in NSCLC microenvironments and can limit the efficacy of immune-checkpoint blockade treatment options such as anti-PD-L1 therapy (Rapoport et al., 2020). We and others have previously reported neutrophil infiltration and low T-cell abundance in *Kras*^*G12D*/+^*;Lkb1*^-/-^ and *Lkb*^-/-^ tumors, which was recapitulated in our ASC^i.v.^ lung colonization model (Koyama et al., 2016, Nagaraj et al., 2017). We have here also reported ASC-specific systemic TAN expansion in KL GEMMs that is also present in i.v. ASC transplanted mice. Underpinning their promise as therapeutic targets, neutrophils have specifically been reported to support the initiation of cancer cell metastasis in solid tumors (Xing et al., 2021, Guan et al., 2021, Wculek and Malanchi, 2015, Tohme et al., 2016, Schneider et al., 2021). Strategies that impair TAN functions may therefore serve as an approach to inhibit metastatic progression, with selective promise for ASC histotype NSCLC. A consequence of neutrophil expansion during inflammation is the formation of stress-associated neutrophil extracellular traps (NET), which have also been shown to regulate both tumor initiation and metastatic processes (Masucci et al., 2020). A strategy to inhibit colorectal cancer growth and metastasis has been to target NETs either directly by local DNase treatment or by inhibition of peptidylarginine deaminase, which is essential for NET formation (Tohme et al., 2016). Another option for targeting neutrophils could be to target chemokines, such as TGF-β1, that have been shown to stimulate the tumor-infiltration by neutrophils (Jayaraman et al., 2018).

In summary, we present a tractable approach to faithfully model NSCLC in a setting that will provide options to investigate metastatic capacities of cells and molecular profiles that dictate responses to targeted therapies. We highlight the unique capacity of KL-ASC, but not KL-AC, tumor cells to effectively re-colonize the lung in recipient i.v. transplanted mice, with histopathological features, oncogenic signaling profiles and previously characterized therapeutic responses maintained. We also show that the TME plays a crucial role in the development of tumors with uniform loss of a benchmark AC molecular marker in subcutaneous tumors independent of the source GEMM lesions. In conclusion, we put forward our lung colonization model as a robust tool to identify improved treatment approaches for NSCLC and solutions to enable the prevention and control of NSCLC tumor metastasis.

## Materials and Methods

### Animal models and tumor initiation in NSCLC GEMMs

All aspects of animal handling and studies were performed by following the guidelines from the Finnish National Board of Animal Experimentation (permit number ESAVI/6365/2019). *Kras^LSL(G12D)/WT^* mice (Jackson et al., 2001) were purchased from The Jackson Laboratory. *Lkb1^fl/fl^* mice (Bardeesy et al., 2002) were received from R. DePinho (MD Anderson, USA). *Kras^LSL(G12D)/WT^* (C57BL/6J background) were bred with *Lkb1^fl/fl^* (F4 ICR;BALB/cByJ;FVB/N background) to generate *Kras^LSL(G12D)/WT^;Lkb1^fl/fl^* (KL) strains (Ji et al., 2007). 3 week old Hsd:AthymicNude-Foxn1nu were purchased from ENVIGO (Huntingdon, UK). To initiate tumor formation, 8-12 week old KL mice (mixed background and C57BL/6 pure background) were intranasally infected with 10^7^ plaque forming units (PFU) AD5-CC10-Cre or 10^9^ PFU Ad5-SPC-Cre (Viral Vector Core Facility, University of Iowa, USA) per mouse in 60 ml total volume of medium (MEM, Sigma) with 9.67 mM CaCl_2_. Mice were sacrificed by cervical dislocation or CO_2_ inhalation upon presentation with labored breathing and weight loss. Both female and male mice were used in similar numbers in all experiments.

### Generation of intravenous and subcutaneous transplantation models

C57Bl/6 *KL* mice were infected with Ad5-CC10-Cre or Ad5-SPC-Cre and were sacrificed upon presentation with labored breathing and weight loss, with median survival of 105 (ASC-bearing mice) and 116 days (AC-bearing mice) post-infection observed. Lungs with tumors were dissected and placed in HBSS. Tumors were isolated from lungs and a tissue reference piece was separated and fixed in 4% paraformaldehyde (MERCK, Kenilwork, NJ, USA) overnight at 4°C. Single cell suspensions were generated from remaining tumor tissue in a digestion mixture with 2 mg/ml Collagenase, 0.3 mg/ml Dispase in HBSS for 30 minutes in 37°C. Tissue pieces were thereafter mechanically disrupted with gentle MACS tube (Miltenyi Biotec) in cold DMEM with 10% FBS, 1% Glutamine and 5 μl/ml DNase using the gentle MACS dissociator. The cells were filtered with a 70 μm cell strainer and dead cell removal was performed on single cell suspensions (Miltenyi Biotec) and live tumor cells were frozen in 90% heat-inactivated fetal bovine serum (HI-FBS) and 10% dimethyl sulfoxide (DMSO). Live tumor cells were revived, washed once with phosphate buffered saline (PBS) and re-suspended in PBS at a concentration of 2 x 10^5^ cells / 100 uμl (for intravenous (i.v.) injection) or 6.6 – 10 x 10^5^ cells / 100 μul (for subcutaneous (s.c.) injection). Single cell suspensions were injected i.v. (tail vein) or s.c. (right flank) into recipient 4-8 week old Hsd:AthymicNude-Foxn1nu or C57BL/6J mice. I.v. injected mice were sacrificed upon presentation with labored breathing and weight loss or reaching 10 weeks post-transplantation. Lungs with tumors were washed with HBSS and fixed in 4% paraformaldehyde overnight at 4°C. Blood was isolated from the saphenous vein immediately following sacrifice, collected into tubes containing 10mM EDTA. Spleens were dissected and bone marrow was flushed from femurs and tibias with a 26 g needle. Spleens were mechanically dissociated (gentleMACS™) white blood cells were isolated from whole blood, bone marrow and spleens by performing red blood cell lysis. White blood cells suspensions from whole blood, spleens and bone marrow were stained with fluorophore-conjugated antibodies for 30 minutes at 4°C and cells were analyzed by flow cytometry (Intellicyt iQUE PLUS screener and Forecyt analysis software). See table S1 for a list of conjugated antibodies and concentrations. Tumor growth in s.c. injected mice were monitored by manual caliper measurements and mice were sacrificed upon i) s.c. tumor reaching 1300 mm^3^; ii) exhibition of inflammation or irritation at tumor site or; iii) reaching 6 months post-injection. Upon sacrifice, s.c. tumors were dissected, washed with HBSS, cut into multiple pieces and fixed in 4% paraformaldehyde overnight at 4°C.

### *In vivo* treatment with afatinib and erlotinib

ASC tumor cells were transplanted into age matched C57BL/6 mice in order to generate tumors. Treatment with afatinib (12.5 mg/kg) or erlotinib (12.5 mg/kg) in 0.5% hydroxyl propyl methyl cellulose and 0.1% Tween 80 in H2O by oral gavage was started 10 days after tumor cell injection. For the analysis of tumor burden at four weeks after injection (Figure 4A-B), mice were treated with afatinib three days a week. In the survival analysis, mice were treated five times a week and sacrificed when their weight dropped more than 20% or health weakened. Mice were sacrificed by cervical dislocation and tumors were collected for IHC analysis.

### Immunohistochemistry (IHC) and histopathology analysis

Formaldehyde fixed tissues were dehydrated in absolute ethanol and isopropanol and embedded in paraffin (FFPE). 4 μm sections were cut and dried overnight at 37°C. FFPE sections were rehydrated, and stained with haematoxylin and eosin (H&E; Merck + Sigma) or IHC using antibodies of interest. Rehydrated tissue sections underwent heat-mediated antigen-epitope retrieval in 10mM Citric Acid buffer pH6 or 10mM Tris/ 1mM EDTA buffer pH9 (Lab Vision™ PT Module, Thermo Fisher Scientific, Waltham, MA, USA), endogenous peroxidase blocking with 0.3% H2O2 (31642, Sigma-Aldrich, St. Louis, MO, USA) for 30 minutes at room temperature and non-specific blocking with 1% bovine serum albumin (BSA; A2153, Sigma-Aldrich) and 10% normal goat serum (NGS; Gibco) for 30 minutes at room temperature. Primary antibodies were diluted in 5% NGS and tissue sections were incubated with primary antibodies for 1-2 hours at room temperature. BrightVision poly-HRP Goat anti-Rabbit (ImmunoLogic, AD Duiven, The Netherlands) was used as the secondary antibody and substrate detection was performed with DAB (Bright DAB, ImmunoLogic). Counterstaining was performed with 10% haematoxylin (Dako) and after dehydration slides were mounted with DPX mounting medium (Merck). Whole slide scans of H&E and IHC-stained lung and tumor sections were acquired with a Panoramic 250 digital slide scanner using a 20x objective (3DHISTECH, Budapest, Hungary). Images of whole slide scans and snapshots were acquired using the online WebMicroscope platform (fimm.webmicroscope.net). See table S1 for a list of IHC antibodies and concentrations.

### *Ex vivo* cell death analysis

Conditionally reprogrammed cultures (CRCs) of ASC and AC tumors isolated from KL-CC10 and KL*-*SPC mice were previously generated and were maintained in co-culture with gamma-irradiated NIH3T3 fibroblasts in F12/DMEM media supplemented with 5% HI-FBS, 5 mg/ml insulin (I2643, Sigma-Aldrich), 24 mg/ml adenine (A2786, Sigma-Aldrich), 10 mM Y-27632 (ALX-270-M0055, Enzo Life Sciences, Farmingdale, NY, USA), 10 ng/ml human recombinant epidermal growth factor (hr-EGF; 354052, BD Pharmigen, San Jose, CA, USA), 0.4 mg/ml hydrocortisone (Sigma), 10 ng/ml cholera toxin (1000B, List Biological Laboratories, Inc., Campbell, CA, USA) and 1% penicillin/streptomycin (15140122, Thermo Fisher Scientific). CRC monocultures were seeded (1.25 x 10^5^ cells per condition), treated with H2O2 (DETAILS, Sigma-Aldrich) and stained with propidium iodide (1 mg/ml, Sigma) prior to analysis with the Intellicyt iQUEÔ HD Screener (Sartorius). Data was analyzed using the Intellicyt Forecyt software.

#### Statistical analysis

Gehan-Breslow-Wilcoxon test was used for the mouse survival curve analysis. One-way Anova test with Tukeys multiple comparisons test was used to test statistical significance in the mouse IHC-based quantifications. To statistically test the rest of the analyses, Student’s two-tailed t test and equal variance was used with following p-values: *p<0.05, **p<0.01, ***p<0.001. All experiments were repeated at least 3 times on different days.

## Author contributions

J.R.D. and E.W.V. conceived the study, and E.W.V. supervised the work. I.A.K.L., J.R.D., A.S.N. and S.T. conducted the experimental design. I.A.K.L., J.R.D., A.S.N., S.T., J.B., N.L., C.S.V., E.A.K. and A.H., performed the experiments. I.A.K.L., J.R.D., A.S.N., S.T., J.B. and E.W.V. conducted data interpretation. I.A.K.L., J.R.D. A.S.N. and E.W.V. wrote the manuscript.

## Acknowledgements

We are grateful to Tomi Mäkelä and Ronald DePinho for the *Lkb^fl/fl^* mice. We thank the FIMM digital microscopy unit for scanning histological slides, and the HiLIFE Laboratory Animal Centre Core Facility at the University of Helsinki for animal husbandry care and support. We thank Kaisa Salmenkivi and Mikko Mäyränpää for histopathology consultation. We thank past and present members of the Verschuren lab for guidance and support. Research was supported by the Liv och Hälsa foundation (E.W.V. & I.A.K.L.), the University of Helsinki Doctoral Programme in Biomedicine (A.S.N, J.B.), the University of Helsinki Integrative Life Science doctoral program (S.S.T.); the Academy of Finland (E.W.V. grant 307111), and the Finnish Cancer Organizations (E.W.V. grant 4706132).

