## Supplementary information for "Development of an adenosquamous carcinoma histopathology-selective lung cancer graft model"

*Shared first authors

Corresponding author: Emmy W. Verschuren (0000-0001-5771-9129)

Keywords: NSCLC, adenosquamous carcinoma, histopathology, metastasis, lung colonization, preclinical model

### Supplementary Figures

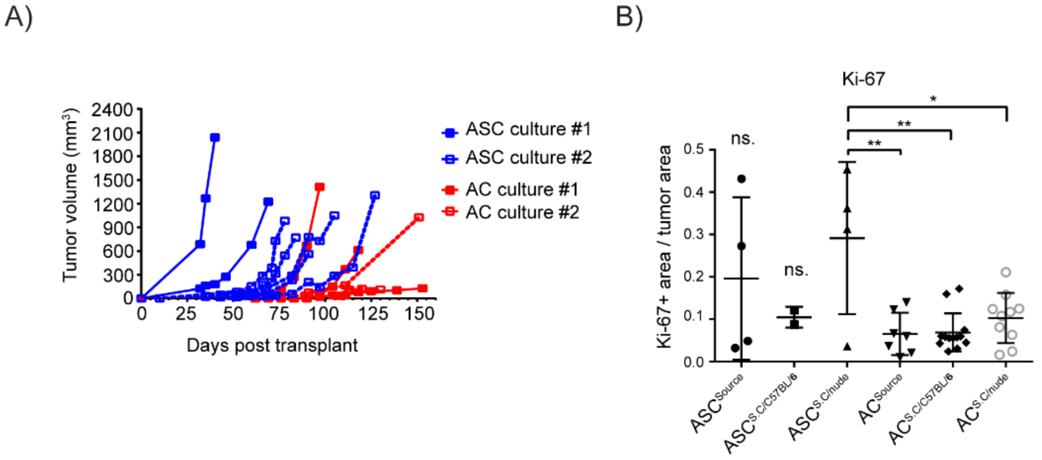


**Figure S1. Host microenvironment-dependent and histopathology-selective tumor establishment subcutaneously. (A)** Subcutaneous ASC or AC tumor growth curves in athymic nude recipient mice using *in vitro* cultured cells. Each line represents one mouse and the tumors were measured weekly by using a caliper. n=7 (ASC), 5 (AC). **(B)** ASC and AC cells were subcutaneously transplanted and post-mortal IHC analysis of Ki-67 was performed (Figure 1 C-D). Quantification of Ki-67 positive area by dividing the positive tumor area with the total tumor area in each tumor separately by using FIJI ImageJ. The graph represents mean+/- SD for each group, and each point represents an individual tumor. Two-tailed unpaired Student's t-test values are *p<0.05, **p<0.01. n=3 (2 for ASC$^{s.c.C57BL/6}$).

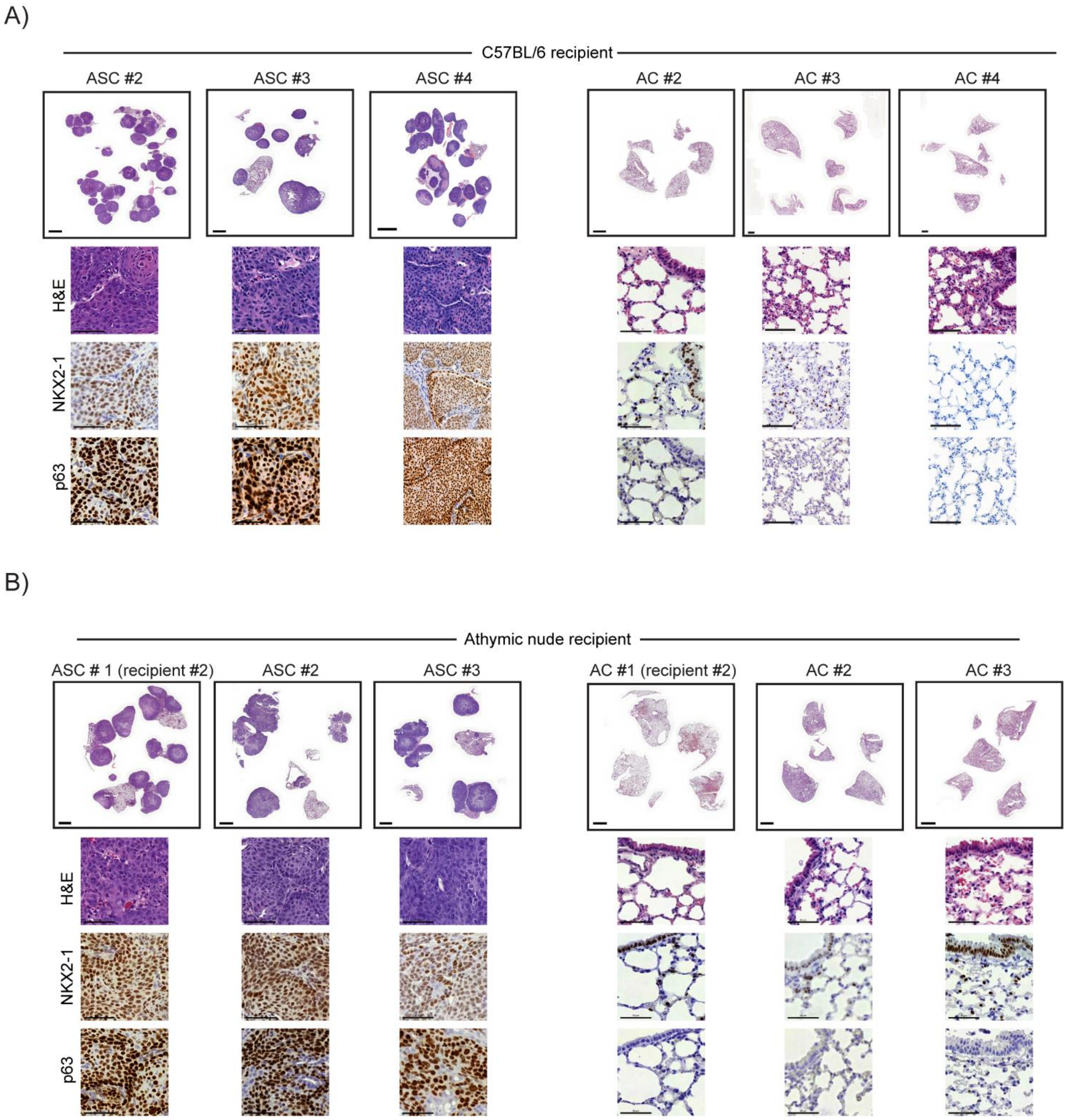


**Figure S2. Host microenvironment-dependent and histopathology-selective tumor establishment intravenously. (A-B)** Biological replicates for Figure 2 with H&E, nkx2-1 and p63, IHC images of C57BL/6 recipient or athymic nude mice upon i.v. transplantation with ASC or AC tumor-derived cells. Scale bar 2 mm (2x) or 50 µm (40x). n=6.

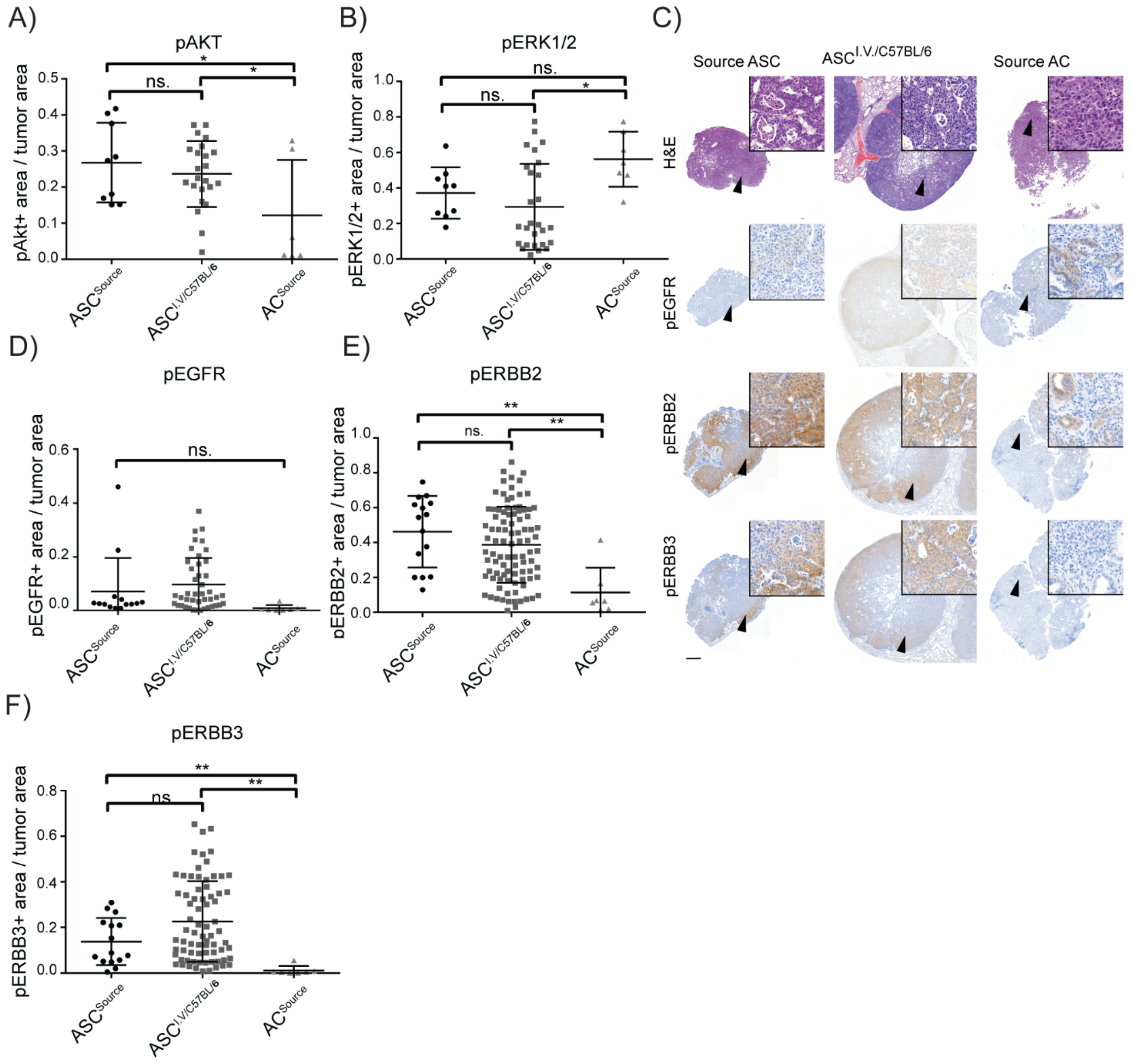


**Figure S3. ASC$^{i.v.}$ cells mimic KL;ASC tumors in showing increased pERBB levels compared to KL;AC mice. (A-B)** Quantification of pAKT and pERK positive area by dividing the positive tumor area with the total tumor area in each tumor separately by using FIJI ImageJ. The graph represents mean+/- SD for each group, and each point represents an individual tumor. Two-tailed unpaired Student's t-test values are *p<0.05. n=5. **(C)** Representative H&E, pEGFR, pERBB2 and pERBB3 IHC images of KL;ASC source, i.v. tumors from C57BL/6 recipient mice upon transplantation with ASC tumor-derived cells and KL;AC source tumors. Scale bar 500 µm (2x) or 20 µm (40x). n=5 **(D-F)** Quantification of pEGFR, pERBB2 and pERBB3 positive area by dividing the positive tumor area with the total tumor area in each tumor separately by using FIJI ImageJ. The graph represents mean +/- SD for each group, and each point represents an individual tumor. Two-tailed unpaired Student's t-test values are *p<0.05, **p<0.01. n=5.

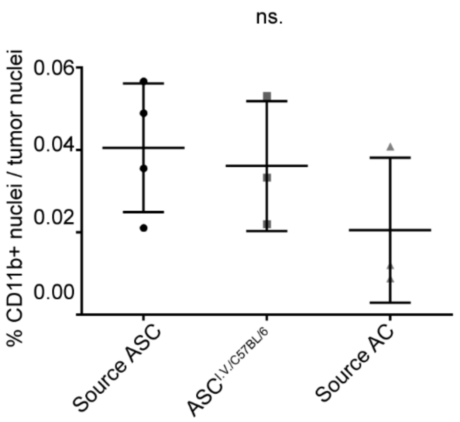


**Figure S4. No differences are measured in the number of CD11b+ neutrophils in KL;ASC source and ASC$^{i.v.C57BL/6}$ tumors.** Quantification of CD11b positive nuclei by dividing the positive tumor nuclei with the total tumor nuclei in each tumor separately by using FIJI ImageJ and CellProfiler. The graph represents mean+/- SD for each group, and each point represents an individual mouse. n=4.

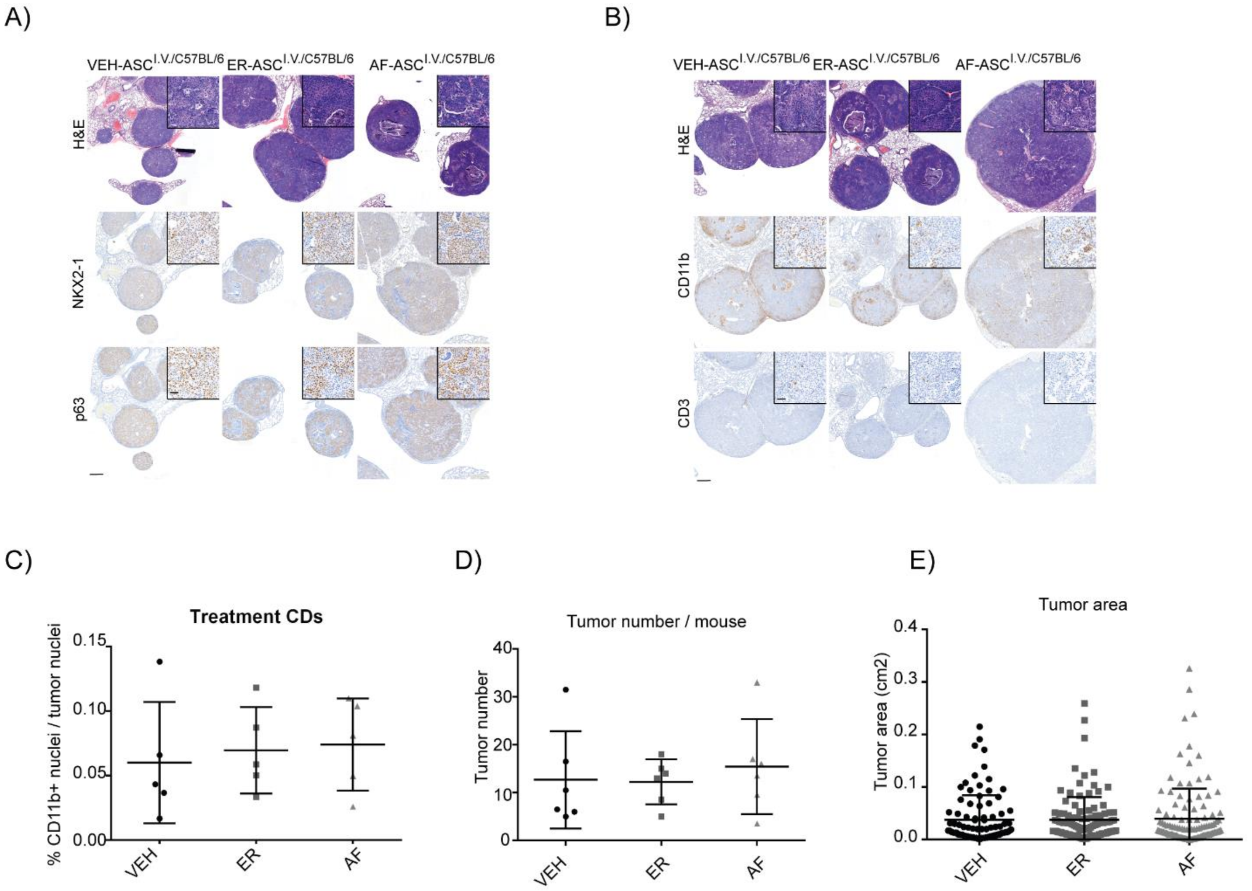


**Figure S5. Therapeutic treatment of $ASC^{i.v.}$ mice does not associate with differences in tumor histotype, immunophenotype or tumor size. (A)** $ASC^{i.v.}$ mice were treated 10 days after intravenous transplantation with vehicle (VEH), erlotinib (ER) or afatinib (AF). Representative H&E, nkx2-1 and p63 IHC images of VEH, ER and AF treated mice. Scale bar 500 µm (2x) or 20 µm (40x). n=5. **(B)** Representative H&E, CD11b and CD3 IHC images of VEH, ER and AF treated mice. Scale bar 500 µm (2x) or 20 µm (40x). n=4-5 **(C)** Quantification of CD11b positive nuclei by dividing the positive tumor nuclei with the total tumor nuclei in each tumor separately by using FIJI ImageJ and CellProfiler. The graph represents mean +/- SD for each group, and each point represents an individual tumor. n=4. **(D)** The tumor number after VEH, ER or AF treatment was derived from the H&E stainings. The graph represents mean +/- SD for each group, and each point represents an individual mouse. n=6. **(E)** The tumor area after VEH, ER or AF treatment was derived from the H&E stainings using Fiji ImageJ. The graph represents mean +/- SD for each group, and each point represents an individual tumor. n=6.

#### Supplementary Table

**Table S1. List of antibodies and their concentrations used for FACS and IHC analysis.**

| Antibody | Company | Code | Contentration | Notes - antigen retrieval |
|---|---|---|---|---|
| p63 | Abcam | ab124762 | 1:10000 | Citric acid pH 6 |
| pAKT | Cell Signaling Tech. | CST4058 | 1:400 | Citric acid pH 6 |
| CD11b | Bio SB | BSB6441 | 1:250 | Citric acid pH 6 |
| CD11b-FITC | Biolegend | 101206 | 1:100 | for FACS |
| CD3 | Abcam | ab5690 | 1:1000 | Citric acid pH 6 |
| CD45- PE | E-Biosciences | 12-0451-82 | 1:200 | for FACS |
| pERK1/2 | Cell Signaling Tech. | CST4370 | 1:1000 | Citric acid pH 6 |
| pEGFR | Cell Signaling Tech. | CST2234 | 1:50 | Tris-EDTA pH 9 |
| pERBB2 | Cell Signaling Tech. | CST2243 | 1:200 | Tris-EDTA pH 9 |
| pERBB3 | Cell Signaling Tech. | CST4791 | 1:200 | Tris-EDTA pH 9 |
| Gr1- APC | Biolegend | 108412 | 1:200 | for FACS |
| KI67 | Thermofisher Scientific | MA-14520 | 1:200 | Tris-EDTA pH 9 |
| NKX2-1 | Abcam | ab133638 | 1:2000 | Citric acid pH 6 |
| Vimentin | Cell Signaling Tech. | CST5741 | 1:500 | Tris-EDTA pH 9 |
